# A novel thermal-visual place learning paradigm for honeybees

**DOI:** 10.1101/2019.12.18.880898

**Authors:** R Scheiner, F Frantzmann, M Jäger, C Helfrich-Förster, D Pauls

## Abstract

Honeybees have fascinating navigational skills and learning capabilities in the field. To decipher the mechanisms underlying place learning in honeybees, we need paradigms to study place learning of individual honeybees under controlled laboratory conditions. Here, we present a novel visual place learning arena for honeybees which relies on high temperatures as aversive stimuli. Honeybees learn to locate a safe spot in an unpleasantly warm arena, relying on a visual panorama. Bees can solve this task very well at a temperature of 46°C, while at temperatures above 48 °C bees die quickly. This new paradigm, which is based on a pioneering work in *Drosophila*, allows us now to investigate thermal-visual place learning of individual honeybees in the laboratory, for example after controlled genetic knockout or pharmacological intervention.

## Results

Despite their minute brain, honeybees have impressive navigational skills, which allow them to locate diverse food sources within a range of several kilometres around their nest and to return fast and direct to their hive (1). A number of experiments suggest that they rely on a cognitive map similar to humans (2, 3). When young bees leave the hive for the first time, they need to acquire information on the local area and perform two days of orientation flights before they begin to forage. If they are displaced during this time, many of them are unable to return to their hive (4). Once they have performed their orientation flights, however, they can be displaced during their outward foraging trip and will still return to their hive using the most direct route, which often involves novel short cuts. Intriguingly, we still know very little about the neuronal mechanisms underlying these complex navigational skills in honeybees.

In order to study visual place learning under controlled laboratory conditions for future analyses on the molecular mechanisms and genetics underlying learning and navigation, we established a thermal visual arena for honeybees, which was adapted from an arena for fruit flies (5). It was inspired by the Morris water maze for rodents and heat mazes for crickets and cockroaches (5–8). In this arena, honeybees learn to escape an unpleasantly hot environment (>44°C) by approaching a safe spot (i.e. a 25°C cool tile) associated with a visual stimulus (Figure 1A-C).

**Figure 1:**
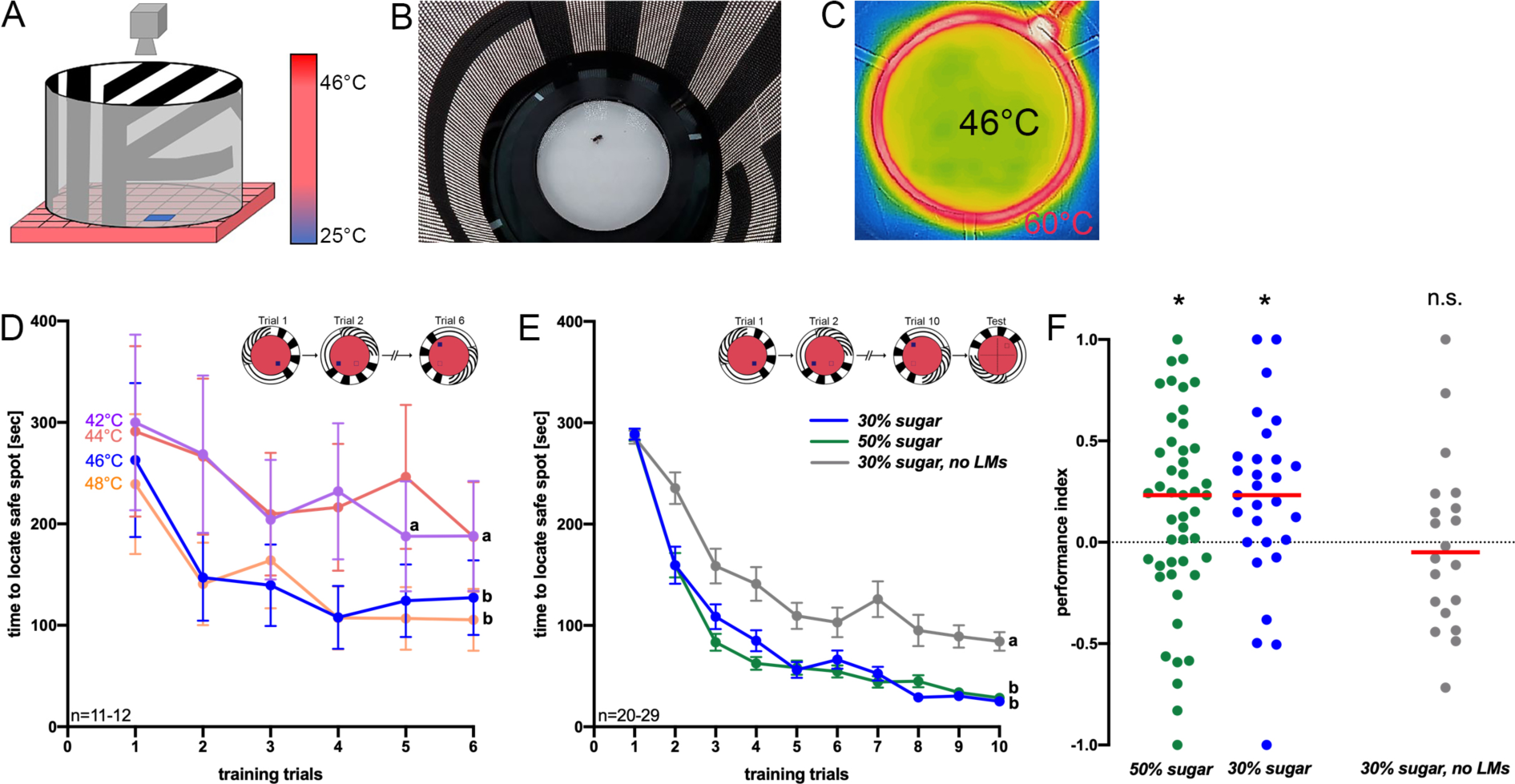
**A.** Schematic picture of the arena for thermal visual place learning in honeybees (adapted from (5)). The ground of the arena is unpleasantly hot and only a small safe spot is (25°C) is offered to walking honeybees. The visual panorama changes with the cold tile. **B**. Photograph of the arena from top. Light emitting diodes illuminate the arena. **C**. Thermal image of the arena: spatial learning experiments were performed using 46°C ground temperature. In addition, a heated ring (~60°C) and a glass lid on top prevent bees from escaping. **D**. Time needed to reach the safe spot at different ground temperatures (42 °C – 48°C) within in six training trials of five min each. Different letters (a,b) indicate significantly different groups. **E**. Times to reach the safe spot using 46°C ground temperature. Of bees which could use landmarks (green and blue graphs; bees of these two groups received different sugar solutions before training) and those which did not have landmarks (grey graph) within ten training trials. The inlet shows examples of the safe spot position. In D and E, mean values and standard errors are shown. **F**. Performance indices of groups trained without landmarks (grey dots) and of those with landmarks (green and blue dots) which differed in the sugar concentration they could feed on prior to training (green: 50% and blue: 30%). (*: P < 0.05, **: P < 0.01; ***: P < 0.001, T test); n.s. = not significantly different from zero.

Honeybees are very sensitive to ambient temperature and maintain a brood nest temperature between 32°C and 36°C, with an optimal temperature of 35°C, to support the appropriate brood development (10). When adult honeybees were tested for their thermotaxis on an aluminium block with a temperature gradient of 28 – 48°C, they preferred temperatures of 34 – 35°C (11). At the individual level, bees avoid temperatures above 44°C and respond with a sting extension to heat stimulations (12). The lethal temperature for honeybees (*Apis mellifera carnica*) is at around 50°C (13). We therefore applied temperatures between 42°C and 50°C to test which temperature is optimal to induce avoidance of honeybees in our arena.

The only spatial cues for the bee in the arena are provided by the surrounding LED panorama (Figure 1A-C). This displays three different stripe patterns with vertical, horizontal and diagonal bars (Figure 1A, 1B; (5)). To assess visual place learning, an individual honeybee is introduced into the arena. During training, we measure the time which the bee needs to reach the safe spot within a time window of five min. A surrounding heated ring (~60°C) and a glass lid prevent the bee from flying off the arena (5). Once the bee has located the safe spot, it normally remains there until the cool tile and the corresponding visual panorama are rotated randomly clockwise or anticlockwise by 90°. Then the bees start searching for the safe spot anew. The bees learn to associate the position of the safe spot relative to that of the visual landmark. Temperatures between 42°C and 50°C (ground temperature) were applied in a first experiment in six trials to investigate at which temperature the bees find the safe spot fastest, following the training trials. At 50°C, five out of seven individuals died after five min in the arena, so that this temperature was abandoned. At the other temperatures (42°C, 44°C, 46°C and 48°C), honeybees became increasingly faster in locating the safe spot with each training session (Figure 1D). The two higher temperatures led to a particularly steep decrease in time to reach the safe spot. For that reason, we selected 46°C (second highest temperature) for further experiments, when asking if bees would use the visual landmarks for orientation. In this experiment, one group of bees (Figure 1E; blue line) could use the T shaped landmark (border area between horizontal and vertical stripes) in the arena, which changed its relative position together with the safe spot (Figure 1E, inset). A second group of bees did not get any landmarks (the screen was set off; Figure 1E, grey graph). Further, we increased the number of training trials to ten, because after six trials not all of the groups tested before appear to have reached an asymptotic performance level. Although both groups showed a significant decrease in time to reach the safe spot (Figure 1E; effect of training trial: F_(6,327)_ = 85.86; P < 0.001, ANOVA RM), the bees which could employ landmarks showed a significantly stronger reduction in the time to reach the safe spot (effect of landmark presence: F_(1,49)_ = 354, P <0.001).

Immediately after training, the bees faced a probe trial without a safe spot to test for visual place memory expression (described in (5)). The landmark changed its position as before. We hypothesized that bees should spend significantly more time in the sector of the arena where the visual landmark suggests the safe spot to be, even though there is no safe spot during testing. The performance index of bees, i.e. the time bees spend in the target sector in comparison to the opposed sector (5), was significantly larger than zero (Figure 1F; T = 5.6, P < 0.001, n = 29), indicating place memory expression. In the absence of landmarks, however, the performance index did not differ from zero (Figure 1F; T = 0.227; P = 0.823, n = 20). The performance indices of both groups differed significantly from each other (T = 3.155; P < 0.01, n = 49).

In many experiments on learning in honeybees (but also in other insects such as *Drosophila* and rodents (14, 15)), the feeding status of the individual has a strong effect on the behavioural response and in particular on learning performance. Accordingly, we hypothesized that place memory expression in bees is dependent on the feeding status, as food deprivation at the individual or colony level is the main driving force for foraging behaviour (16, 17). We therefore asked if learning in a thermal-visual arena may be affected by the sugar concentration which the bees had access to before training. Bees fed with either 30% or 50% *ad libitum* sugar solution before training displayed a significant decrease in time to reach the safe spot (effect of training trial: F_(9,684)_ = 209,32; P < 0.001, ANOVA RM), while the time to locate the safe spot of both groups did not differ statistically (factor feeding status: F_(1,76)_ = 0.68; P = 0.411, ANOVA RM). During the test, the performance index of both groups differed significantly from zero (30 %: T = 2.51, n = 29, P < 0.05; 50 %: T = 2.18, n = 47, P < 0.05) suggesting place memory expression. The performance index of the two groups receiving different sugar solutions did not differ significantly (T = 0.50; P > 0.05).

Taken together, our results show that honeybees can rely on visual landmarks to locate and learn a safe spot position in an otherwise hot arena. Further, the bee’s performance in this non-food reinforced task is independent of the feeding regime, at least for the selected standard sugar solutions (30% and 50%) given to the bees before the day of training. Our paradigm allows us now to study navigational skills of individual honeybees under controlled laboratory conditions, enabling manipulations and intervention with neuronal signalling pathways to understand the neuronal mechanisms underlying visual navigation and learning.

## Methods

### Animals

All experiments were performed with honeybees (*Apis mellifera carnica*) from queen-right colonies maintained at the departmental apiary of Würzburg University. Colonies have been treated against *Varroa destructor* regularly with a sufficient time interval to experiments. Experimental animals were collected from the hive entrance and wings were cut on the same day of the experiment or the day before, depending on the experiment, to prevent the bees from flying about. Bees were kept overnight in an incubator for 24h at 28°C and 60% humidity. Bees could feed *ad libitum* from either 30% or 50% sugar solution until testing.

### Behavioral experiments

Spatial learning experiments were performed in a visual heat maze arena, which was described for the first time by Ofstad et al. 2011. For a more detailed description see (5).

### Statistics

The effect of training trial and of treatment on the time needed to reach the safe spot was compared between different groups using repeated measurement analysis of variance (ANOVA RM, factor training trial or factor treatment, SPSS, IBM). To test whether the learning index differed from zero, one sample T tests were performed. The learning indices of two different groups were compared using independent T tests. All tests were two-tailed.

